# Newly identified parasitic nematode beta-tubulin alleles confer resistance to benzimidazoles

**DOI:** 10.1101/2021.07.26.453836

**Authors:** Clayton M. Dilks, Emily J. Koury, Claire M. Buchanan, Erik C. Andersen

## Abstract

Infections by parasitic nematodes cause large health and economic burdens worldwide. We use anthelmintic drugs to reduce these infections. However, resistance to anthelmintic drugs is extremely common and increasing worldwide. It is essential to understand the mechanisms of resistance to slow its spread. Recently, four new parasitic nematode beta-tubulin alleles have been identified in benzimidazole (BZ) resistant parasite populations: E198I, E198K, E198T, and E198stop. These alleles have not been tested for the ability to confer resistance or for any effects that they might have on organismal fitness. We introduced these four new alleles into the sensitive *C. elegans* laboratory-adapted N2 strain and exposed these genome-edited strains to both albendazole and fenbendazole. We found that all four alleles conferred resistance to both BZ drugs. Additionally, we tested for fitness consequences in both control and albendazole conditions over seven generations in competitive fitness assays. We found that none of the edited alleles had deleterious effects on fitness in control conditions and that all four alleles conferred strong and equivalent fitness benefits in BZ drug conditions. Because it is unknown if previously validated alleles confer a dominant or recessive BZ resistance phenotype, we tested the phenotypes caused by five of these alleles and found that none of them conferred a dominant BZ resistance phenotype. Accurate measurements of resistance, fitness effects, and dominance caused by the resistance alleles allow for the generation of better models of population dynamics and facilitate control practices that maximize the efficacy of this critical anthelmintic drug class.

**Highlights:**

- Four newly identified parasitic nematode beta-tubulin alleles confer benzimidazole resistance
- The four newly identified alleles do not confer deleterious fitness consequences
- Five beta-tubulin alleles confer recessive benzimidazole resistance

**Graphical abstract:** 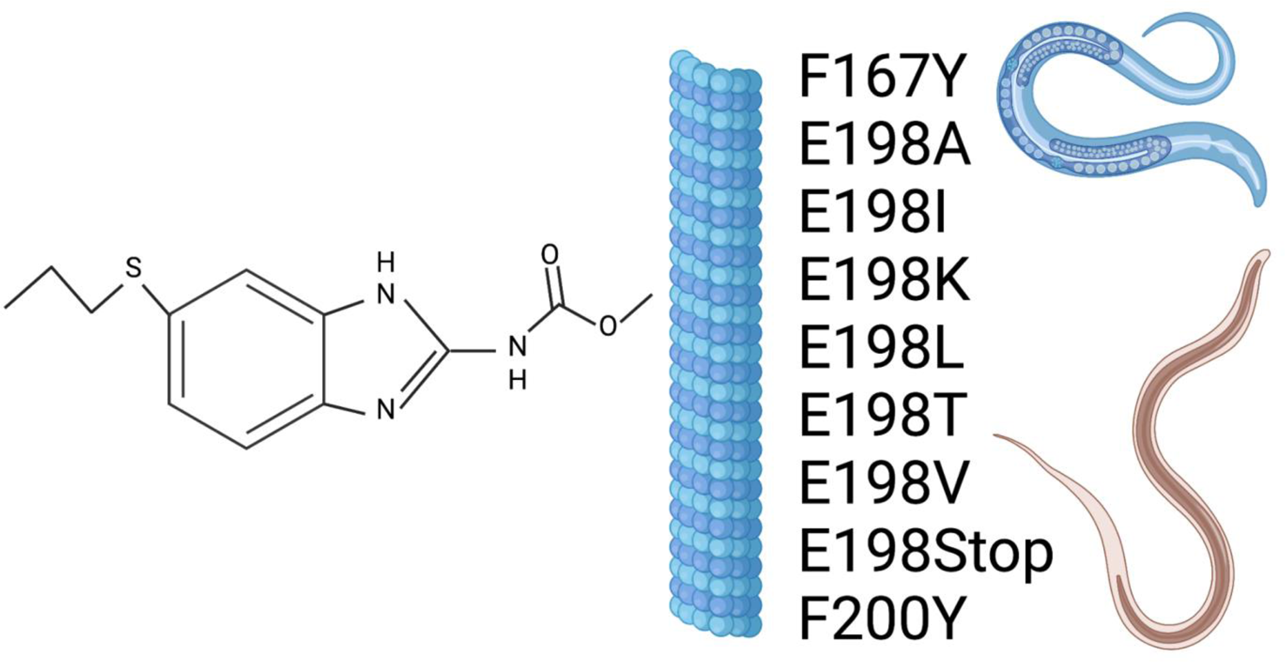

## 1. Introduction

Parasitic nematode infections impact the lives and livelihoods of billions of people each year (Kaplan and Vidyashankar, 2012). These infections can cause a wide range of diseases and cause severe effects, including anemia, malnutrition, and increased susceptibilities to other diseases (Stepek et al., 2006). In addition to humans, many species of livestock are regularly infected by parasites, which cause severe disease in animal health. Researchers estimate that hundreds of millions of dollars in revenue are lost yearly because of parasitic nematode infections of livestock (Kahn and Woodgate, 2012).

Anthelmintic drugs are the major course of treatment against parasitic nematode infections (Abongwa et al., 2017), including four major drug classes: benzimidazoles (BZs), macrocyclic lactones (MLs), nicotinic acetylcholine receptor agonists (NAChAs), and amino- acetonitrile derivatives (AADs). BZs were introduced first and quickly became widely used as the primary treatment for infected livestock, regardless of burden. This treatment strategy causes selection for resistance over a short period of time (Roos et al., 1995), so resistant populations of parasites were found three years after the introduction of BZ compounds (Theodorides et al., 1970). Resistant populations have been selected in disparate sites throughout the world and are now extremely common to the extent that nearly all surveyed parasitic nematode populations have BZ resistance <u>(Howell et al. 2008; Crook et al. 2016)</u>.

To prevent the spread of resistance, proper herd control and pasture management using techniques such as refugia will help lower the levels of resistant parasites in a population (Hodgkinson et al., 2019). The refugia technique maintains a population of susceptible parasites in hosts separated from the main herd and rotates this population back into the main herd when resistance alleles are present at high frequencies (Hodgkinson et al., 2019). For the refugia technique to be effective, a comprehensive catalog of resistance alleles and the effects of those alleles must be defined to enable proper monitoring and planning of refugia pasture rotations (Hodgkinson et al., 2019; Wit et al., 2021).

Resistance to BZ compounds is the most thoroughly understood of all the drug classes. Early studies focused on BZ resistance in fungi found that BZs target beta-tubulin (Hastie and Georgopoulos, 1971; Sheir-Neiss et al., 1978). Later research using the free-living nematode *Caenorhabditis elegans* found that BZ resistance can be conferred by loss of a gene that encodes beta-tubulin (Driscoll et al., 1989). Using these findings, researchers began to identify alleles associated with resistance in beta-tubulin genes in parasite populations, such as F200Y (Roos et al., 1990; Kwa et al., 1993, 1994). However, the presence of beta-tubulin alleles in resistant populations does not mean that these alleles confer resistance. In order to show a causal connection between an allele and resistance, experiments must address both sufficiency and necessity of an allele in BZ resistance. Sufficiency has been shown for beta-tubulin in BZ resistance using *C. elegans* extrachromosomal array experiments (Kwa et al., 1995). However, necessity requires a single allele to be introduced into a susceptible background and for it to confer resistance. This proof has recently been obtained for five of the parasitic nematode beta- tubulin alleles: F176Y, E198A, E198L, E198V, and F200Y <u>(Hahnel et al., 2018; Kitchen et al.,</u> <u>2019; Dilks et al., 2020)</u>. Additionally, the ability of an allele to confer resistance is only one factor used to understand how an allele impacts a parasite population. An allele that confers a high level of resistance but also causes an organism to be less fit will not spread in a population as fast as one that confers resistance without the fitness consequences. Parasite populations must balance the fitness effects of alleles and the selective pressure from BZ drugs. For example, a resistance-conferring allele that negatively affects fitness when drug pressure is absent can be controlled by a prolonged period without drug treatment or by treatment with another drug class. The dominance of a resistance phenotype also impacts how an allele will spread in a parasite population. If an allele arises that confers a dominant resistance phenotype, then this allele will quickly increase in frequency across the population because it has an immediate impact on drug resistance. Conversely, if an allele arises that confers a recessive resistance phenotype, the allele will spread more slowly because it will not be selected unless homozygous (Cornell et al., 2003).

As sampling of parasite populations increased over the last few decades, an extensive catalog of putative beta-tubulin resistance alleles have been discovered in parasites: F167Y, E198A, E198I, E198K, E198L, E198T, E198V, E198stop, and F200Y (Silvestre and Cabaret, 2002; Ghisi et al., 2007; von Samson-Himmelstjerna et al., 2009; Redman et al., 2015; Avramenko et al., 2019; Mohammedsalih et al., 2020). The quantitative levels of BZ resistance and fitness effects of five of these alleles (F167Y, E198A, E198L, E198V, and F200Y) have been determined previously (Dilks et al., 2020; Hahnel et al., 2018; Kitchen et al., 2019). In this study, we tested the effects of four newly discovered parasite beta-tubulin alleles on BZ resistance (E198I, E198K, E198T, and E198stop). Using CRISPR-Cas9 genome-editing, we introduced these alleles into the BZ-susceptible laboratory-adapted strain, N2, and performed high-throughput assays to test for resistance to two BZs, albendazole and fenbendazole. We found that all four alleles conferred BZ resistance similar to a strain with a *ben-1* deletion allele. In addition to these BZ-response assays, we performed competitive fitness assays to identify any fitness effects conferred by these alleles. We found that all edited alleles did not confer deleterious fitness effects in control conditions. Additionally, we found that all edited alleles conferred similar survival advantages in BZ drug conditions. Finally, we performed a quantitative dominance assay to test the dominance relationship of five alleles, F167Y, E198A, E198V, and F200Y, and found that they each did not confer a dominant BZ resistance phenotype. Validation, levels of resistance, and dominance effects of these alleles allow for accurate modeling of how resistant alleles will impact and spread throughout parasite populations.

## 2. Methods

### 2.1 *C. elegans* strains

Animals were grown at 20°C on a modified nematode growth media (NGMA) plates that contained 1% agar and 0.7% agarose with OP50 bacteria (Andersen et al., 2014). For all assays, strains were grown for three generations to alleviate multigenerational effects of starvation (Andersen et al., 2015). The previously tested BZ-resistant *ben-1* deletion strain, ECA882 *ben-1(ean64)*, was used as a resistant control in all assays (Hahnel et al., 2018). The wild-type strain with a barcoded *dpy-10* gene, PTM229 *dpy-10(kah81)*, was used in competitive fitness assays (Zhao et al., 2018). All strains used in this manuscript (Supplemental Table 1) were generated in the N2 background with modifications introduced using CRISPR-Cas9 genome editing as previously described (Hahnel et al., 2018; Dilks et al., 2020) and below.

### 2.2 CRISPR-Cas9 genome editing

Genome editing was performed using a co-CRISPR strategy in the N2 genetic background, as previously described (Kim et al., 2014; Hahnel et al., 2018; Dilks et al., 2020). We designed sgRNAs to target the *ben-1* and *dpy-10* loci to create the E198I, E198K, E198T, and E198stop allele-replacement strains. The online analysis platform Benchling (www.benchling.com) was used for all guide RNA (sgRNA) designs. The sgRNAs were ordered from Synthego (Redwood City, CA) and injected at 5 µM and 1 µM for the *ben-1* and *dpy-10* guides, respectively. Single-stranded oligodeoxynucleotides (ssODN) templates were used for homology-directed repair (HDR) of both targeted loci. These ssODNs were ordered as ultramers (IDT, Coralville, IA) and injected at 6 µM for the *ben-1* repair and 0.5 µM for the *dpy- 10* repair. Purified Cas9 protein (QB3 Macrolab, Berkeley, CA) was injected at 5 µM. All reagents were incubated at room temperature for one hour before injection into the germlines of young adult hermaphrodite animals. Injected animals were singled to new 6 cm NGMA plates 18 hours post-injection. After two days, F1 progeny were screened for the Rol phenotype, and Rol individuals were singled to new NGMA plates. The ssODNs contained a silent edit to the PAM site and another silent edit recognized by the *Cla*I restriction enzyme. Following PCR of the targeted region, the amplicon was incubated with the *Cla*I enzyme (New England Biolabs, Ipswich, MA). Successful edits were identified by an altered restriction pattern of wild-type amplicons compared to the edited amplicons. F2 non-Rol offspring of successfully edited parents were singled to new NGMA plates. The edited genomic regions from F2 individuals were then Sanger sequenced to ensure homozygosity of the edit. Two independent edits of each allele were generated to control for off-target effects that can occur during the CRISPR- Cas9 genome-editing process. All oligonucleotides in this study are available upon request (Supplemental Table 2).

### 2.3 High-throughput fitness assays

High-throughput fitness assays were performed as previously described (Zdraljevic et al., 2017; Brady et al., 2019; Dilks et al., 2020; Evans and Andersen, 2020). In short, a 0.5 cm^3^ NGMA chunk was removed from an NGMA plate that contained starved individuals and placed onto a fresh NGMA plate. After 48 hours, gravid hermaphrodites were placed into a bleach solution on a new 6 cm NGMA plate to clean each strain. The following day, five L1 larvae were transferred to a 6 cm NGMA plate and allowed to develop and reproduce for five days. After five days, the offspring had reached the L4 stage, and five L4 animals were placed on a new 6 cm NGMA plate and allowed to develop for four days. Following this growth, strains were bleached in 15 mL conical tubes to generate a large pool of unhatched embryos. Three independent bleaches were performed to control for variability introduced by the bleach synchronization process. The embryos were then diluted to approximately one embryo per µL in K medium (51 mM NaCl, 32 mM KCl, 3 mM CaCl2, and 3 mM MgSO4 in distilled water) (Boyd et al., 2012). The embryo suspension was then placed into 96-well plates at approximately 50 embryos per well and allowed to hatch overnight. The next day, lyophilized bacterial lysate (*E. coli* HB101 strain (García-González et al., 2017)) at a concentration of 5 mg/mL were fed to the population of animals. In addition to the bacterial lysate, albendazole in DMSO, fenbendazole in DMSO, or DMSO alone were added at the same time. A final concentration of 1% DMSO was maintained for all conditions. Both albendazole and fenbendazole were used at a final concentration of 30 µM as described previously (Dilks et al., 2020). After 48 hours of growth, animals were scored using the COPAS BIOSORT (Union Biometrica, Holliston MA) after treatment with 50 mM sodium azide to straighten the animals (Zdraljevic et al., 2017; Hahnel et al., 2018; Brady et al., 2019; Dilks et al., 2020; Evans and Andersen, 2020). Animal optical density normalized by animal length (EXT) was calculated for each animal. Benzimidazole treatment delays the development of susceptible animals and reduces their optical density compared to control animals, as previously described (Dilks et al., 2020). The median optical density of the population within each well was used to quantify benzimidazole responses. Growth trait processing and analyses were performed using the R(4.0.3) package *easysorter* (Shimko and Andersen, 2014) with the v3_assay = TRUE option in the *sumplate* function. Analyses were performed as described previously (Dilks et al., 2020). All normalizations were performed on a strain-specific basis to account for any differences present in the starting populations. All data and scripts for this analysis are available at https://github.com/AndersenLab/2021_ben1resistance_CMD.

### 2.4 Competition assays

We used an established pairwise competition assay to assess competitive fitness (Hahnel et al., 2018; Zhao et al., 2018; Dilks et al., 2020). The competitive fitness of each strain was determined by comparing allele frequencies of the test strain with a wild-type control strain. The PTM229 strain contains a silent mutation in the *dpy-10* locus to differentiate the strains using oligonucleotide probes that specifically bind to the barcoded PTM229 version or the wild-type version (Zhao et al., 2018). Ten fourth larval stage individuals from both the PTM229 (wild-type *ben-1* strain) and *ben-1* edited strains were placed on a single 6 cm NGMA plate containing either DMSO (control) or 1.25 µM albendazole in DMSO (Hahnel et al., 2018;Dilks et al., 2020). Ten independent competitions were set up for each condition and strain combination. The N2 and ECA882 (*ben-1(ean64)*) strains were included as controls for sensitivity and resistance, respectively, because they have previously been characterized (Hahnel et al., 2018; Dilks et al., 2020). Plates were grown for one week until starvation. A 0.5 cm^3^ piece of agar was then transferred to a new NGMA plate of the same condition to seed the next generation. We then washed the remaining animals off the plates into a 1.7 mL microfuge tube with M9 buffer, spun the tubes in a centrifuge, and removed as much liquid as possible from the pellet. Samples were then stored at -80°C. DNA was extracted using the DNeasy Blood & Tissue kit (Qiagen 69506). All competitions were performed for seven generations, and DNA was collected from generations one, three, five, and seven.

We quantified the allele frequencies of each strain as previously described (Zhao et al., 2018). Briefly, a droplet digital PCR approach using TaqMan probes (Applied Biosciences) was used. Extracted DNA was digested using *EcoR*I for 30 minutes at 37°C, purified with Zymo DNA cleanup kit (D4064), and diluted to 1 ng/µL. With previously designed TaqMan probes (Zhao et al., 2018), we performed droplet digital PCR using a Bio-Rad QX200 device with standard probe absolute quantification settings. One TaqMan probe binds selectively to the wild-type *dpy-10* allele and the other binds to the PTM229 *dpy-10* allele. The relative allele frequencies of each competition were calculated using the QuantaSoft software with default settings. A one-locus generic selection model was used to calculate the relative fitness of each allele (Zhao et al., 2018).

### 2.5 Dominance assay

We used a modified version of the previously described high-throughput fitness assay (Brady et al., 2019; Evans et al., 2020) to assess the dominance relationships for the resistance conferred by six *ben-1* alleles (Δ*ben-1*, F167Y, E198A, E198L, E198V, and F200Y) and the wild-type allele. We set up a cross between a sensitive strain and a resistant strain to test this dominance relationship. To differentiate cross progeny from single-parent offspring, we used an N2-derivative strain that constitutively expresses GFP (EG8072). The strains harboring edited *ben-1* alleles were grown as described for the high-throughput assay. Thirty L4 hermaphrodites of each strain were placed on a NGMA plate with 60 males of EG8072 and allowed to mate for 48 hours. After 48 hours, the animals on these plates were bleached in 15 mL conical tubes and embryos aliquoted into 96-well plates at approximately one embryo per µL of K-medium and left to hatch overnight. The next day, the hatched L1 larvae were fed bacterial lysate along with 30 µM albendazole in DMSO or DMSO at a final concentration of 1% for both conditions. After 48 hours of growth, the populations were scored using the COPAS BIOSORT as previously described in the high-throughput fitness assay.

After scoring the individuals with the COPAS BIOSORT, we performed an analysis to differentiate cross-progeny from self-progeny in each well. All cross-progeny express GFP because the male parent harbored a transgene array that expresses GFP. To identify GFP-expressing individuals, we used the *easysorter* package (Shimko and Andersen, 2014), which outputs the green fluorescence of each individual scored using the COPAS BIOSORT. The green value is then normalized by the length of the individual to obtain the norm.green trait. The distribution of norm.green across the population was bimodal, so we used the local minimum of the distribution to set a cutoff to determine if an animal expressed GFP. This cross design gave us the ability to measure the differences in responses between heterozygous individuals and homozygous individuals at the *ben-1* locus from the same cultures. Traits for testing resistance were analyzed using the same method previously described in high-throughput fitness assays.

### 2.6 Statistical analysis

All statistical tests and comparisons were performed in *R* version 4.0.3 (Core Team and Others, 2013) using the *Rstatix* package. The *Rstatix tukeyHSD* function was used on an ANOVA model (formula = *phenotype ∼ strain*) to calculate differences among strains BZ response.

### 2.7 Research Data

Supplementary Table 1 contains a list of all strains and genotypes, along with primer and guide RNA sequences. Supplemental Table 2 includes all oligonucleotide sequences used in the study. All scripts and data for this manuscript are available at https://github.com/AndersenLab/2021_ben1resistance_CMD.

## 3. Results

### 3.1 The E198I, E198K, E198T, and E198stop beta-tubulin alleles confer resistance to benzimidazoles in *C. elegans*

We performed CRISPR-Cas9 genome editing to introduce each parasitic nematode beta-tubulin allele into the *C. elegans ben-1* gene in the N2 genetic background. For each candidate resistance allele, two independent edited strains were created. This experimental design made it possible to test only the effect of this single amino-acid change on the responses to albendazole and fenbendazole. We then performed high-throughput assays to test if the E198I, E198K, E198T, and E198stop alleles conferred resistance to albendazole and fenbendazole. We included the *C. elegans* N2 (laboratory wild-type) strain and a strain with the *ben-1* locus deleted as controls for susceptible and resistant strains, respectively. The assay included 21 replicates per strain with 35-50 animals per replicate in each drug and control condition. We performed a high-throughput assay that measured optical density as a response to BZ treatment as previously described (Dilks et al., 2020) because this trait is a proxy for developmental rate. Higher regressed optical density values correspond to increased resistance to the tested BZs (larger animals that developed further), and lower values correspond to increased sensitivity to the tested BZs (smaller animals that were developmentally delayed). All tested parasitic nematode beta-tubulin alleles showed significant increases in resistance compared to the susceptible laboratory strain and similar resistance levels to the resistant *ben-1* deletion strain (**Fig. 1A, B, Supplemental Fig. 1**). These results indicate that all four newly identified alleles conferred similar levels of BZ resistance as the previously tested strains, F167Y, E198A, E198L, E198V, and F200Y (Hahnel et al., 2018; Dilks et al., 2020). In control conditions, the strains containing the E198T and E198stop alleles grew significantly faster compared to the strains containing the wild-type, E198I, and E198K alleles (**Supplemental Fig. 2**). This increased growth compared to the wild-type strain was surprising because we did not expect to find any differences among strains in control conditions. To more directly test this result suggesting increased fitness in control conditions, we tested these alleles in a pairwise competition assay against the laboratory wild-type strain.

**Figure 1:**
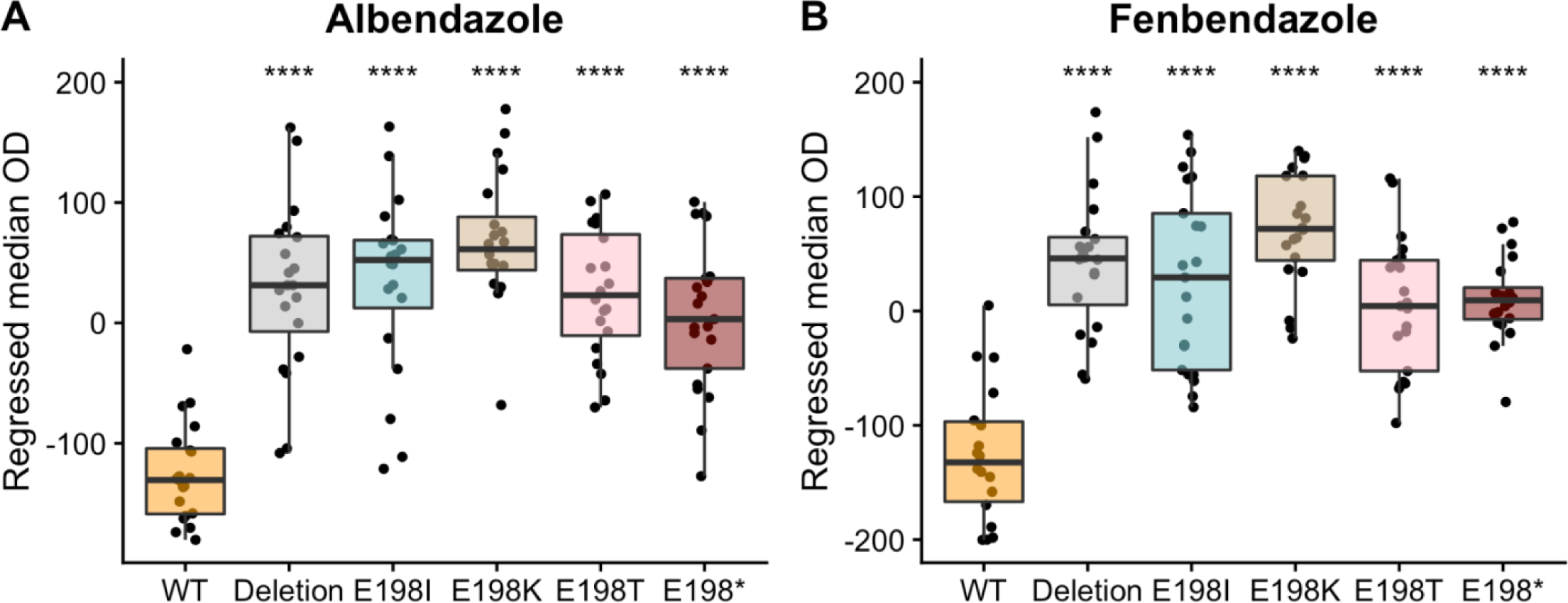
Drug-response assays for the E198I, E198K, E198T, and E198stop parasite beta-tubulin alleles. Regressed median optical density (median.EXT) values from populations of nematodes grown in either 30 µM albendazole (A) or 30 µM fenbendazole (B) are shown on the y-axis. Each point represents the regressed median optical density value of a well containing approximately 35- 50 animals. Data are shown as Tukey box plots with the median as a solid horizontal line, and the top and bottom of the box representing the 75th and 25th quartiles, respectively. The top whisker is extended to the maximum point that is within 1.5 interquartile range from the 75th quartile. The bottom whisker is extended to the minimum point that is within 1.5 interquartile range from the 25th quartile. Significant differences between the wild-type strain and all other alleles are shown as asterisks above the data from each strain (p < 0.0001 = ****, Tukey HSD).

### 3.2 The E198I, E198K, E198T, and E198stop alleles do not cause negative fitness consequences in control conditions

We performed a seven-generation competitive fitness assay to measure the fitness of strains with edited *ben-1* alleles compared to the wild-type *ben-1* allele. We performed these competitions in both DMSO and ABZ because the level of fitness in both conditions is important for parasite control models. This assay is more sensitive than our one generation high-throughput growth assay because small changes in fitness can be amplified over multiple generations. If a strain is more fit than the unedited wild-type control strain, then that strain will increase in frequency in each generation. By contrast, if the unedited wild-type control strain is detected at higher frequency across the generations, then the edited allele likely confers a fitness disadvantage in those conditions, as observed previously with the E198V *ben-1* allele (Dilks et al., 2020). We also competed the N2 strain and a strain with *ben-1* deleted with the wild-type barcoded strain to use as controls for sensitive and resistant ABZ strains, as we have done previously (Hahnel et al., 2018; Dilks et al., 2020).

We found that the wild-type strain had no difference in fitness compared to the barcoded wild-type strain in both control and drug conditions (**Fig 2**). Additionally, the *ben-1* deletion strain had no fitness consequences in control conditions but a strong competitive advantage in drug conditions. Next, we investigated the effects of the genome-edited beta-tubulin alleles. In control conditions, all genome-edited *ben-1* strains had levels of fitness statistically indistinguishable from the control unedited wild-type strain (**Fig 2A,B**). In ABZ conditions, all edited *ben-1* alleles increased in frequency across the seven generations (**Fig 2C**) and had significantly higher competitive fitness than the unedited wild-type strain (**Fig 2D**). These results suggest that all of these alleles cause similar levels of resistance with little to no fitness costs.

**Figure 2:**
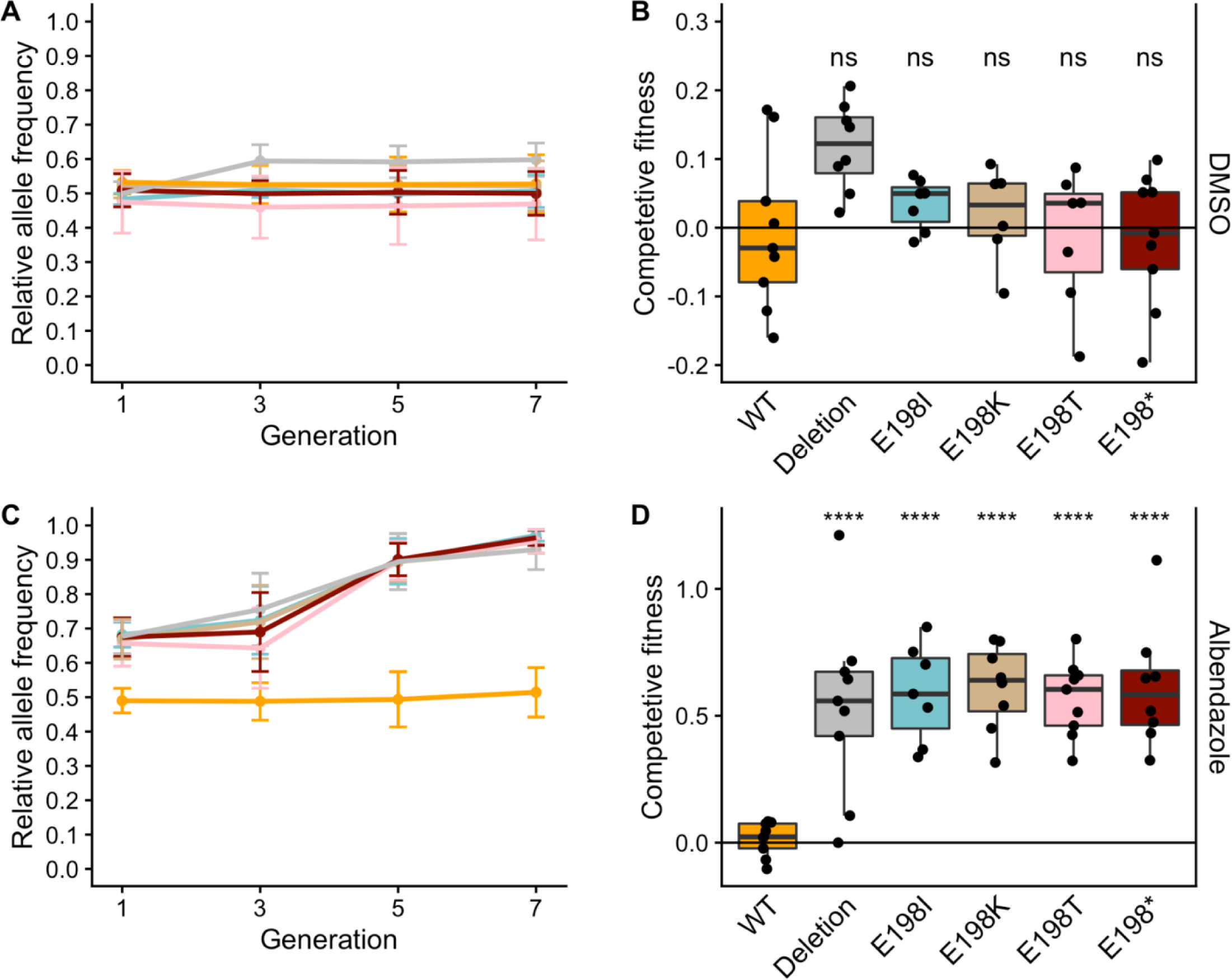
Competitive fitness assay across seven generations in both DMSO and ABZ. (A) A barcoded N2 wild-type strain was competed with strains harboring different *ben-1* alleles in DMSO control conditions. The generation is shown on the x-axis, and the relative allele frequency of the genome-edited *ben-1* allele is shown on the y-axis. (B) The log2-transformed competitive fitness of each allele is plotted. The allele tested is shown on the x-axis, and the competitive fitness is shown on the y-axis. Each point represents a biological replicate of that competition experiment. Data are shown as Tukey box plots with the median as a solid horizontal line, and the top and bottom of the box representing the 75th and 25th quartiles, respectively. The top whisker is extended to the maximum point that is within 1.5 interquartile range from the 75th quartile. The bottom whisker is extended to the minimum point that is within 1.5 interquartile range from the 25th quartile. Significant differences between the wild-type strain and all other alleles are shown as asterisks above the data from each strain (p > 0.05 = ns, p < 0.0001 = ****, Tukey HSD). (C) A barcoded N2 wild-type strain was competed with strains harboring different *ben-1* alleles in 1.25 µM ABZ. (D) The log2-transformed competitive fitness of each allele is plotted. Each point represents one biological replicate of the competition assay. Data are plotted as in (B).

### 3.3 Five parasitic nematode beta-tubulin alleles (F167Y, E198A, E198V, E198L, and F200Y) do not confer dominant BZ resistance phenotypes

We performed a high-throughput assay to quantitatively test dominance of the resistance phenotype conferred by five beta-tubulin alleles (F167Y, E198A, E198V, E198L, and F200Y). We used a modified version of the previously described high-throughput assay (see Methods) in which the animals tested were offspring from a cross between a wild-type male expressing GFP and a non-GFP-expressing hermaphrodite (**Fig. 3A**). Therefore, cross progeny expressed GFP and were heterozygous at the *ben-1* locus (and all other loci). To identify cross progeny, we looked at the distribution of normalized GFP expression in the experiment (**Fig. 3B**). A clear bimodal distribution was observed as expected for a mixed population of GFP-expressing and non-GFP-expressing individuals. We then calculated the local minimum in the bimodal distribution and used that value as a threshold to designate an individual as GFP-positive or GFP-negative. We then measured the difference between heterozygous and homozygous responses to albendazole to determine if the BZ-response caused by these alleles was dominant or recessive.

**Figure 3:**
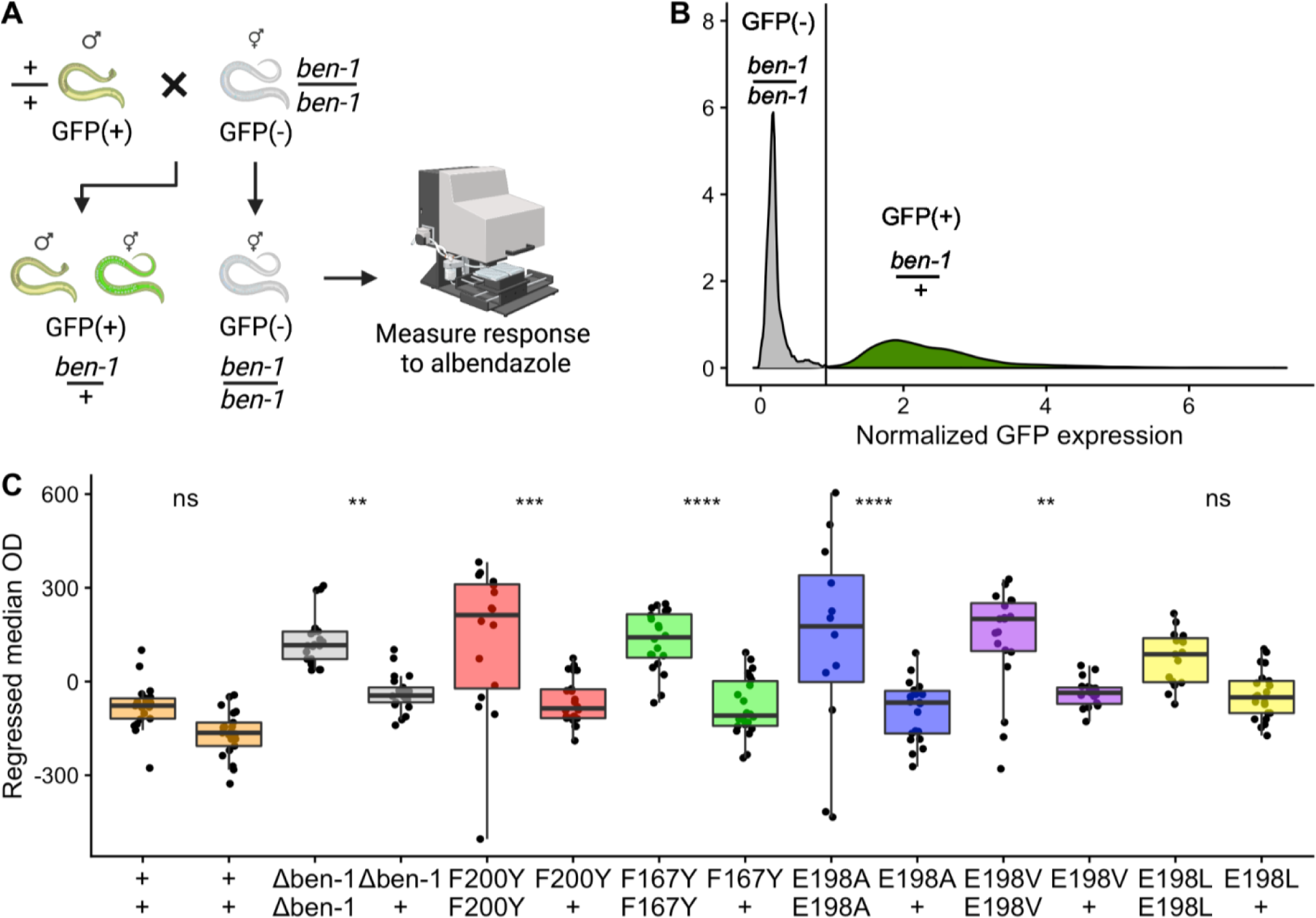
Quantitative dominance tests of parasite beta-tubulin alleles in albendazole. A) A graphical representation of the dominance assay experiment is shown. GFP-positive (GFP(+)) wild-type males and GFP-negative (GFP(-)) hermaphrodites harboring different *ben- 1* alleles were crossed for 48 hours. Following mating, two categories of animals were present, GFP(+) and GFP(-). GFP(+) individuals were heterozygous for the *ben-1* locus, and GFP(-) individuals were homozygous at the *ben-1* locus. These populations of animals were then exposed to albendazole and scored using the COPAS BIOSORT. This illustration was generated on biorender.com. (B) The x-axis represents the total GFP expression of each animal normalized by animal length, and the y-axis represents the distribution of the population of animals. The vertical line denotes the threshold used to differentiate animals expressing GFP and animals not expressing GFP (see Methods). The distribution is filled based on if the individuals in that part of the distribution were categorized as expressing GFP (green) or not (gray). The genotypes of animals in each category are shown above the distributions. (C) The x-axis shows the genotype at the *ben-1* locus for animals in the dominance assay. Regressed median optical density values of responses to 30 µM albendazole are shown on the y-axis. Each point represents the regressed median optical density value from a population of 35-50 animals in a single well. Data are shown as Tukey box plots with the median as a solid horizontal line, the top and bottom of the box representing the 75th and 25th quartiles, respectively. The top whisker is extended to the maximum point that is within 1.5 interquartile range from the 75th quartile. The bottom whisker is extended to the minimum point that is within 1.5 interquartile range from the 25th quartile. Significant differences between the homozygous genotype and the heterozygous genotype are shown above each of the tested alleles (p < 0.01 = **, p<0.001 = ***, p < 0.0001 = ****, Tukey HSD).

The *ben-1* deletion, F167Y, E198A, E198V, and F200Y alleles showed significant differences between the homozygous and heterozygous individuals for 22 replicate populations in albendazole (**Fig. 3C**). The wild-type strain and E198L did not show a significant difference between heterozygous and homozygous individuals. It appeared that the E198L allele was similar to the other parasitic nematode beta-tubulin alleles but the difference between heterozygous animals and homozygous individuals was not significant (p = 0.225, Tukey HSD) likely because of experimental noise. In control conditions, both the heterozygous individuals and the self-progeny showed no significant differences in growth (**Supplemental Fig. 3**). These results indicated that at least five and maybe all six of these alleles did not confer a dominant BZ resistance phenotype at this concentration of albendazole in laboratory conditions.

## 4. Discussion

### 4.1 Newly identified parasitic nematode alleles confer resistance to BZs in *C. elegans*

Recent work has shown that the F167Y, E198A, E198L, E198V, and F200Y alleles confer resistance to BZs (Hahnel et al., 2018; Kitchen et al., 2019; Dilks et al., 2020). Because of increased surveillance and sequencing of BZ resistant parasitic nematode populations, more beta-tubulin alleles have been identified recently in *H. contortus* isotype-1: E198I, E198K, E198T, and E198stop (Mohammedsalih et al., 2020). The identification of new alleles is important for proper parasite control measures because it allows more accurate surveillance of resistance alleles in populations. Once identified, these alleles need to be validated because not all changes to a gene sequence will have functional effects and many alleles might be correlated or linked with other causal loci. Misclassification of resistance alleles could negatively impact control measures where genetic tests optimize the use of refugia. Although technology for CRISPR-Cas9 genome editing is still being developed for parasitic nematode species, *C. elegans* is easily manipulated using this technology and provides an excellent platform for validation of newly discovered parasitic nematode beta-tubulin alleles in BZ resistance (Hahnel et al., 2018; Kitchen et al., 2019; Dilks et al., 2020). Here, we introduced the E198I, E198K, E198T, and E198stop alleles into a defined *C. elegans* genetic background and showed that each allele confers a high level of resistance similar to loss of the *ben-1* locus.

We also performed pairwise competition assays to test for fitness effects of these newly identified alleles. The high-throughput assay and competition assay have different advantages and disadvantages, as previously discussed (Dilks et al., 2020). In short, the high-throughput assay allows testing of a large number of strains and quickly can confirm if alleles confer resistance to BZs. The pairwise competition assay is more sensitive to small differences in growth in control (DMSO) or BZ conditions. For example, previous work using this assay identified a small fitness cost associated with the E198V allele compared to the wild-type allele (Dilks et al., 2020). This difference in fitness was not observed over a single generation (as is used in the high-throughput fitness assays) so both high-throughput resistance and pairwise competition assays are required to test both of the effects caused by beta-tubulin alleles. This study, along with previous research (Hahnel et al., 2018; Kitchen et al., 2019; Dilks et al., 2020), provides a complete catalog of the effects of all identified parasitic nematode beta-tubulin alleles.

Using current technologies, we can rapidly validate quantitative levels of BZ resistance and measure fitness effects after discovery of variants from parasitic nematode populations. The F200Y allele was discovered almost thirty years ago (Kwa et al., 1994) but was not experimentally validated until recently (Hahnel et al., 2018). By contrast, the E198K allele was published in 2020 (Mohammedsalih et al., 2020), and we confirmed the resistance of this allele in this study. The speed of validation is important for parasitic nematode control because it gives researchers the ability to implement resistance mitigation techniques like refugia and drug-class rotation to lower the levels of resistance parasitic nematode populations.

### 4.2 The new beta-tubulin alleles confer fitness advantages in BZ conditions and no deleterious fitness effects in control conditions

We tested the fitness of all newly identified beta-tubulin alleles in highly sensitive competitive fitness assays. We found that no strains had a statistically significant difference in competitive fitness from the unedited N2 strain in control conditions (**Fig 2A,B**). The *ben-1* deletion allele was slightly more fit compared to the E198stop allele (p=0.0319, TukeyHSD), but this effect was not significant when compared to any other allele. Therefore, we believe that this difference is not biologically relevant and likely represents assay-to-assay variation. Our results here and published previously (Dilks et al., 2020) show that, among all beta-tubulin alleles found in parasitic nematodes, only the E198V allele causes deleterious fitness effects in control conditions. All of the alleles confer quantitatively equal levels of BZ resistance (Dilks et al., 2020) and (**Fig 2C,D**). Parasitic nematodes with beta-tubulin mutations must balance the fitness consequences in conditions without BZ compounds with resistance in the presence of these compounds.

Without obvious deleterious effects on fitness, parasitic nematode beta-tubulin resistance alleles should increase in frequency whenever BZ compounds are given to infected hosts. Our results indicate the beat-tubulin alleles confer quantitatively equal levels of resistance. It remains unanswered why only a subset of beta-tubulin positions are found altered in parasite populations and why these resistance alleles vary in frequency across different parasite populations. These observations are seemingly in contrast to the resistance and fitness studies performed here using *C. elegans* in a laboratory setting. Future experiments should alter the beta-tubulin repertoire of *C. elegans* to more closely resemble parasitic nematode genomes. This change in “dosage” might alter the dynamics of resistance. Additionally, the levels of quantitative BZ resistance and fitness effects caused by these alleles might be environment-specific. Resistance assays and pairwise competition experiments at different temperatures, using different bacterial foods, and other environmental alterations could reveal more about how these alleles act in the context of parasitic nematodes.

### 4.3 Parasitic nematode beta-tubulin alleles do not confer a dominant resistance phenotype in *C. elegans*

Because the dominance of BZ resistance caused by beta-tubulin alleles can impact control of parasite populations, we tested the dominance caused by six beta-tubulin alleles (Δ*ben-1*, F167Y, E198A, E198L, E198V, and F200Y). We found that five of the six tested beta-tubulin alleles conferred a recessive albendazole-response phenotype. The sixth allele (E198L) followed a similar trend as the other alleles, but the difference was not statistically significant. However, when population summary statistics are not used and individual animal responses are used for each allele, E198L responds similarly to the other parasitic nematode beta-tubulin alleles (**Supplemental Fig. 4**). The recessive BZ phenotypes suggest that these alleles will not spread as quickly in populations as they would if they caused dominant BZ resistance. When a resistance allele that confers recessive BZ resistance appears in a parasite population, it will be heterozygous and not be selected in BZ conditions. It will need to be homozygous for selection to act on it, so this allele must be in two individuals of the opposite sex and they must mate. This finding suggests that refugia methods, where sensitive parasite populations are mixed into resistant populations, could be an effective method to reduce the spread of resistance alleles because it would ensure that many alleles are found in the heterozygous state. However, resistance alleles in *H. contortus* populations seem to persist long after BZ drug selection is removed (van Wyk et al., 1997), so our results on dominance and validated resistance will be needed to more effectively track the effects of these alleles in parasite populations.

Our findings are inconsistent with earlier *C. elegans* studies where mutations in *ben-1* conferred dominant resistance phenotypes (Driscoll et al., 1989). This difference could be explained by assay temperature. The previous study found that the resistance phenotype was recessive at 15°C but the phenotype was dominant at 25°C (Driscoll et al., 1989). We performed our study at 20°C. It is possible that we could observe a dominant BZ resistance phenotype if we repeated our assays at 25°C. Additionally, the previous study was only performed on males (Driscoll et al., 1989), suggesting that sex might also play a role in dominance.

This interaction between dominance and temperature could suggest that in tropical environments, where the average temperature is warmer than temperate climates, these same beta-tubulin alleles could confer dominant resistance and spread faster. Some previous studies in parasitic nematodes found that BZ resistance was incompletely dominant (Jambre et al., 1979; Martin et al., 1988; Dobson et al., 1996), so more experiments in varying environmental conditions are required to understand BZ resistance more clearly. Parasitic nematodes interact with multiple environments throughout diverse life cycles. For example in *H. contortus*, the temperature within the host organism is much different from the temperature the larvae will experience on the ground in the pasture. Besides temperature, other factors such as salinity, food availability, and the surrounding microbial environment could all be investigated for potential effects on the dominance of the BZ resistance trait.

### 4.4 Future Directions

Our findings in this study can be directly applied to parasite control practices in parasitic nematodes. In order for the refugia method to be successful, a susceptible population of parasites must be maintained. It is imperative to have constant monitoring for the appearance of resistance alleles to keep frequencies at levels where sensitive refugia populations can drive resistance down. Additionally, farms are not closed systems; the livestock population is altered as individuals move among different farms. With the information we now have with regards to BZ resistance alleles, we can screen livestock before they are moved between farms to be sure resistance alleles are not spread between populations. This study can help inform future parasite control strategies and slow the spread of resistance to BZs.

## Acknowledgements

We want to thank Katie Evans and Janneke Wit for their feedback and comments on this manuscript. C.M.D. was supported by the Biotechnology Training Program at Northwestern University (T32 GM008449). E.C.A. was funded by the National Institutes of Health NIAID grant R21AI121836. This study used data supplied by Wormbase, the *Caenorhabditis* Genetics Center (P40 OD010440), and the *Caenorhabditis elegans* Natural Diversity Resource (NSF CSBR 1930382). The funding sources had no impact on the design of this study. We would like to thank biorender.com for the generation of figure 3A and the graphical abstract.

---

**Supplemental figure 1:**
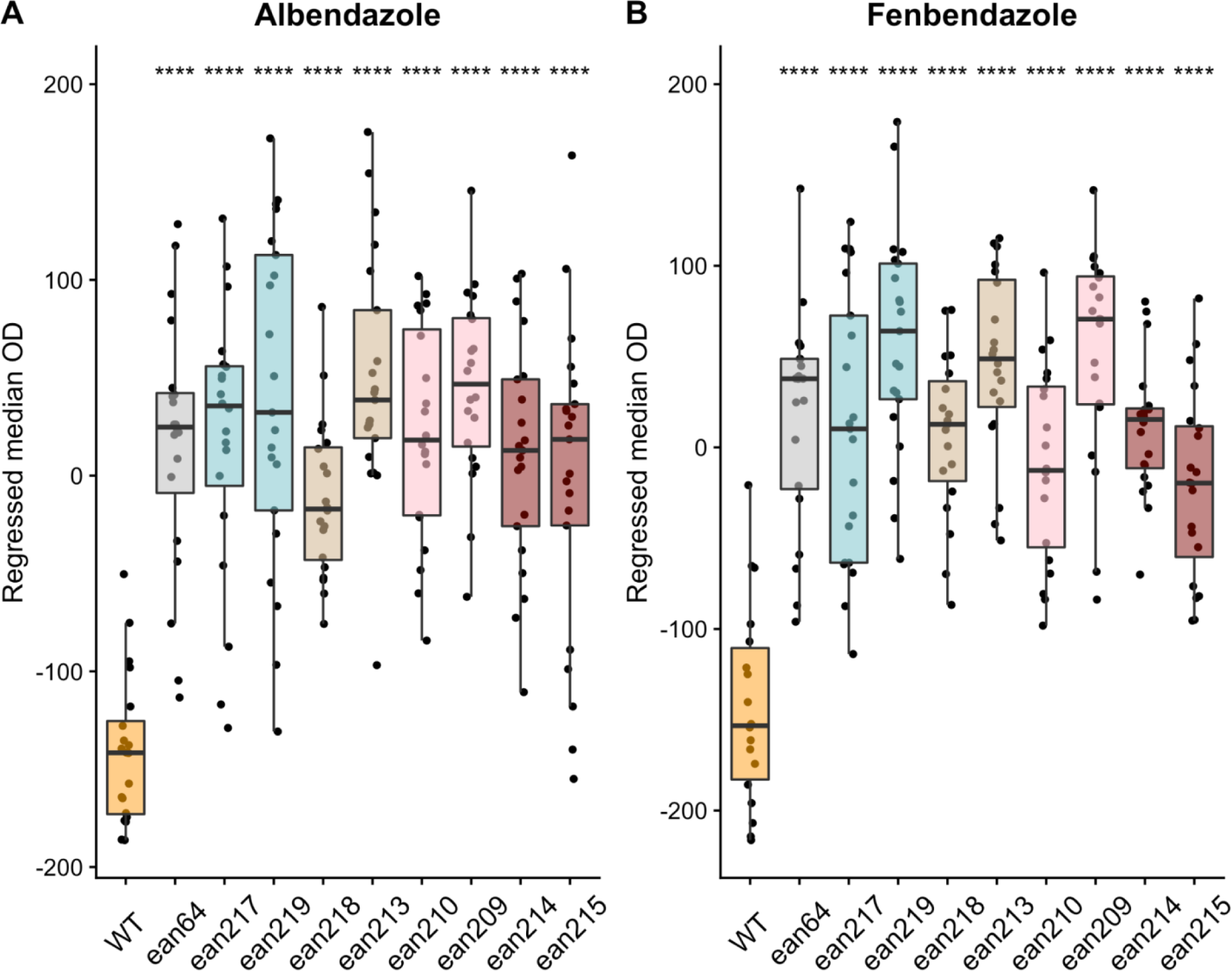
Independent alleles measured in high-throughput assays. Regressed median optical density (median.EXT) values from populations of nematodes grown in either 30 µM albendazole (A) or 30 µM fenbendazole (B) are shown on the y-axis. The x-axis denotes the *ben-1* allele designation of each tested strain (*ean64 =* Δ*ben-1*, *ean217* = E198I, *ean219* = E198I, *ean218* = E198K, *ean213* = E198K, *ean210* = E198T, *ean209* = E198T, *ean214* = E198*, *ean215* = E198*). Each point represents the regressed median optical density value from approximately 35-50 animals in a single well. Data are shown as Tukey box plots with the median as a solid horizontal line, the top and bottom of the box representing the 75th and 25th quartiles, respectively. The top whisker is extended to the maximum point that is within 1.5 interquartile range from the 75th quartile. The bottom whisker is extended to the minimum point that is within 1.5 interquartile range from the 25th quartile. Significant differences between the wild-type strain and all other strains are shown as asterisks above the data from each strain (p < 0.0001 = ****, Tukey HSD).

**Supplemental figure 2:**
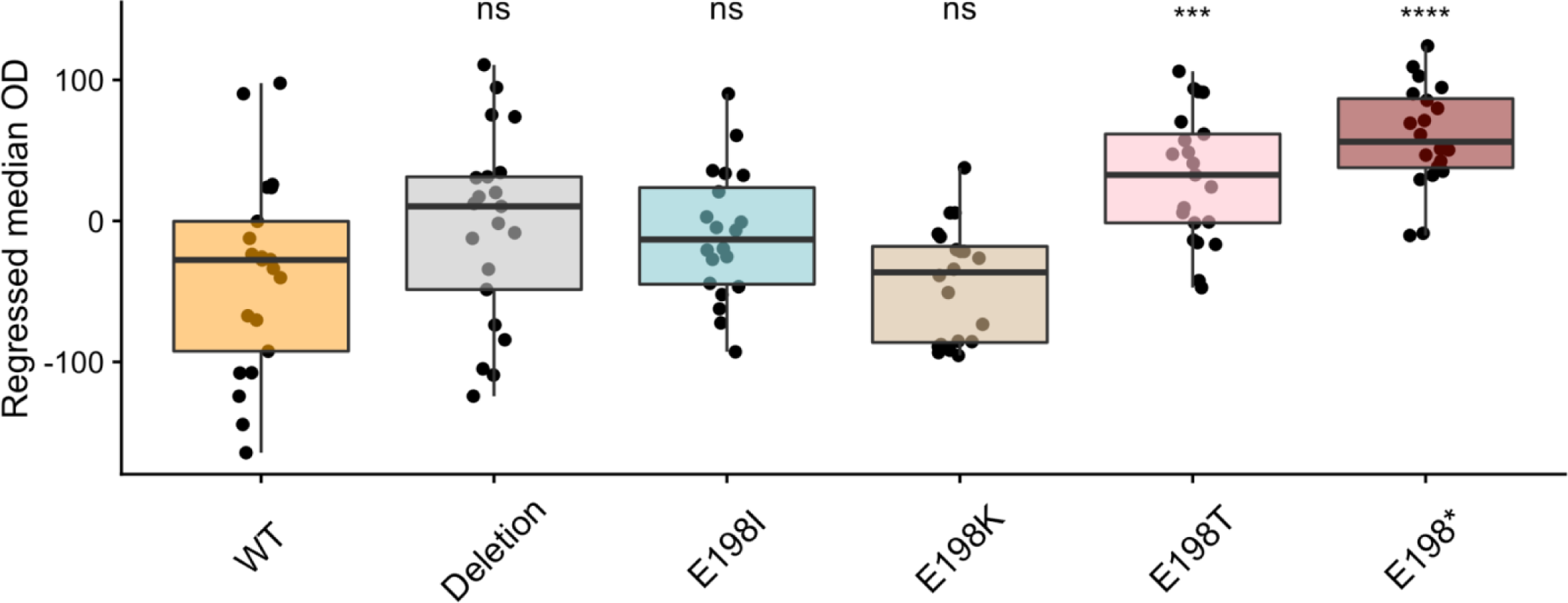
Control response for the E198I, E198K, E198T, and E198* parasite beta-tubulin alleles. Regressed median optical density (median.EXT) values from populations of nematodes grown in DMSO are shown on the y-axis. Each point represents the regressed median optical density value from a well containing approximately 35-50 animals. Data are shown as Tukey box plots with the median as a solid horizontal line, the top and bottom of the box representing the 75th and 25th quartiles, respectively. The top whisker is extended to the maximum point that is within 1.5 interquartile range from the 75th quartile. The bottom whisker is extended to the minimum point that is within 1.5 interquartile range from the 25th quartile. Significant differences between the wild-type strain and all other strains are shown as asterisks above the data from each strain (p < 0.001 = ***, p < 0.0001 = ****, Tukey HSD).

**Supplemental figure 3:**
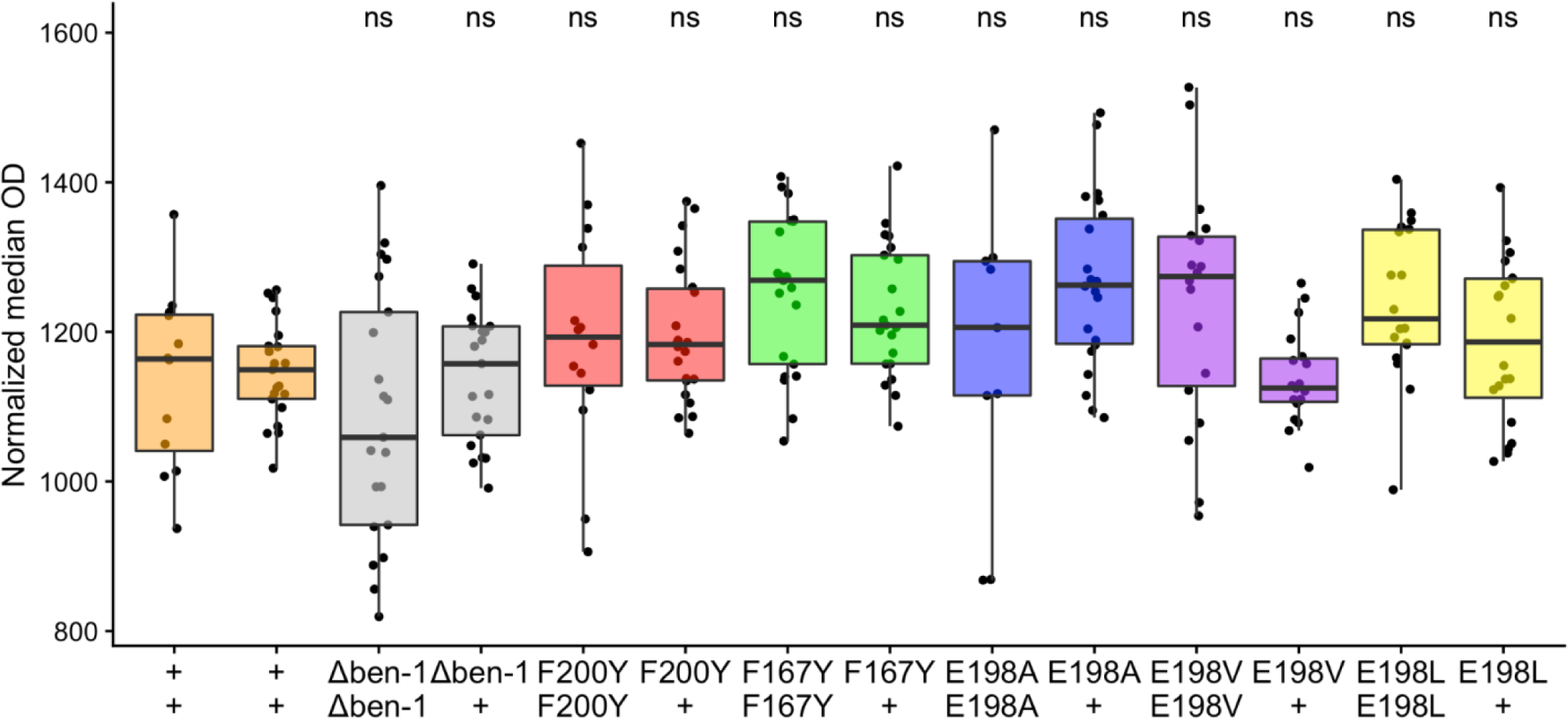
Alleles measured in DMSO conditions following cross. Normalized optical density values of strains in control conditions are shown on the y-axis. The x-axis shows the genotype for *ben-1*. Each point represents the normalized median optical density value from a population of 35-50 animals in a single well. Data are shown as Tukey box plots with the median as a solid horizontal line, the top and bottom of the box representing the 75th and 25th quartiles, respectively. The top whisker is extended to the maximum point that is within 1.5 interquartile range from the 75th quartile. The bottom whisker is extended to the minimum point that is within 1.5 interquartile range from the 25th quartile. No significant differences between N2 and any other strains was identified (p > 0.05, Tukey HSD).

**Supplemental figure 4:**
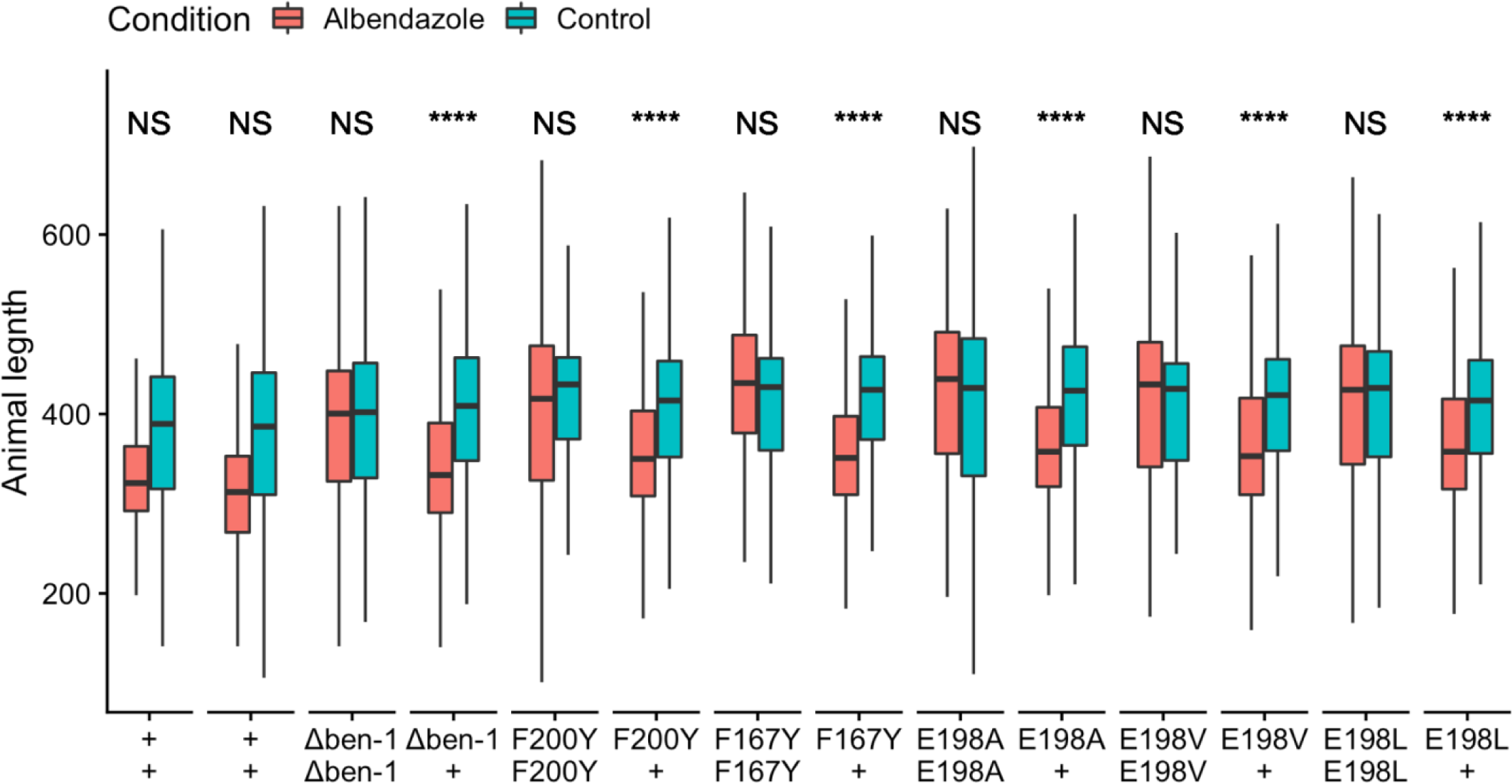
Raw length data of each genotype. The x-axis denotes the genotypes tested. Animal length measurement values (time of flight) are shown on the y-axis. Measurements consist of all individual animals within each group. Significance between control and albendazole conditions are shown above each genotype response ( NS = p > 0.05, **** = p < 0.0001, TukeyHSD)

